# ABCF1 regulates dsDNA-induced immune responses in human airway epithelial cells

**DOI:** 10.1101/2020.02.08.940023

**Authors:** Quynh T. Cao, Jennifer A. Aguiar, Benjamin J-M Tremblay, Nadin Abbas, Nicholas Tiessen, Spencer Revill, Nima Makhdami, Anmar Ayoub, Gerard Cox, Kjetil Ask, Andrew C. Doxey, Jeremy A. Hirota

## Abstract

**Background:** The airway epithelium represents a critical component of the human lung that helps orchestrate defences against respiratory tract viral infections, which are responsible for more than 2.5 million deaths/year globally. Innate immune activities of the airway epithelium rely Toll-like receptors (TLRs), nucleotide binding and leucine-rich-repeat pyrin domain containing (NLRP) receptors, and cytosolic nucleic acid sensors. ATP Binding Cassette (ABC) transporters are ubiquitous across all three domains of life – Archaea, Bacteria, and Eukarya – and expressed in the human airway epithelium. ABCF1, a unique ABC family member that lacks a transmembrane domain, has been defined as a cytosolic nucleic acid sensor that regulates CXCL10, interferon-β expression, and downstream type I interferon responses. We tested the hypothesis that ABCF1 functions as a dsDNA nucleic acid sensor in human airway epithelial cells important in regulating antiviral responses.

**Methods:** Expression and localization experiments were performed using *in situ* hybridization and immunohistochemistry in human lung tissue from healthy subjects, while confirmatory transcript and protein expression was performed in human airway epithelial cells. Functional experiments were performed with siRNA methods in human airway epithelial cells. Complementary transcriptomic analyses were performed to explore the contributions of ABCF1 to gene expression patterns.

**Results:** Using archived human lung and human airway epithelial cells, we confirm expression of ABCF1 gene and protein expression in these tissue samples, with a role for mediating CXCL10 production in response to dsDNA viral mimic challenge. Although, ABCF1 knockdown was associated with an attenuation of select genes involved in the antiviral responses, Gene Ontology analyses revealed a greater interaction of ABCF1 with TLR signaling suggesting a multifactorial role for ABCF1 in innate immunity in human airway epithelial cells.

**Conclusion:** ABCF1 is a candidate cytosolic nucleic acid sensor and modulator of TLR signaling that is expressed at gene and protein levels in human airway epithelial cells. The precise level where ABCF1 protein functions to modulate immune responses to pathogens remains to be determined but is anticipated to involve IRF-3 and CXCL10 production.

## Introduction

The human lung functions at the interface of the external and internal environments and is exposed to over 10,000 litres of air each day from normal respiration. The airway epithelium represents a critical component of the human lung that helps orchestrate defences against inhaled noxious substances that may include air pollution, allergens, bacteria, and viral insults(1–3). To manage these continuous insults, the airway epithelium has evolved to be a multi-functional barrier tissue with mechanical and immunological impedances, manifested through the mucociliary ladder, protein-protein junctions, and innate immune processes. A dominant exposure important in both healthy individuals and those with underlying chronic respiratory diseases are viral infections. Collectively, respiratory tract viral infections are responsible for more than 2.5 million deaths/year globally and represent an economic burden on health care systems for all demographics(4). In individuals with underlying chronic airway disease, respiratory tract viral infections increase frequency and severity of disease exacerbations, hospitalizations, and contribute to morbidity and mortality(5–9). Understanding the mechanisms governing respiratory tract viral infections and host defence is essential for the future development of treatments aimed at minimizing the morbidity and mortality of these pathogens.

Innate immune activities of the airway epithelium rely on accurate sensing of the external environment. The threat posed by viruses that infect the respiratory mucosa is countered by the airway epithelium expressing functional Toll-like receptors (TLRs), nucleotide binding and leucine-rich-repeat pyrin domain containing (NLRP) receptors, and cytosolic nucleic acid sensors that are able to rapidly detect exposures and provide host defence(1–3, 10–12). Antiviral sensing mechanisms in the respiratory mucosa enable responses to influenza A, respiratory syncytial virus, rhinovirus, and human parainfluenza virus; all single stranded RNA viruses(13). dsDNA viruses are also relevant lung infections, with adenovirus capable of inducing influenza like symptoms in healthy subjects and associated with chronic respiratory disease exacerbations (8, 14–16). Like RNA viruses, adenovirus is able to infect airway epithelium followed by replication, which leads to a variety of innate immune defences able to sense viral nucleic acids and proteins(14, 17, 18). Vaccinia virus is another dsDNA virus that is able to infect airway epithelium and has been explored for capacity to genetically engineer the virus for transgene delivery, vaccination strategies, and studying Variola virus infections (19–23). Exploring how the airway epithelium responds to viruses may provide new strategies for controlling infections, optimizing transgene delivery, and vaccination strategies relevant in lung health and disease.

ATP Binding Cassette (ABC) transporters are ubiquitous across all three domains of life – Archaea, Bacteria, and Eukarya(24). In humans, the 49 ABC transporters are classified according to structure and function, resulting in 7 families. ABC transporters with clear involvement in lung health and disease include ABCA3 and ABCC7 (better known as cystic fibrosis transmembrane conductance regulator – CFTR), responsible for surfactant production and ion transport, respectively (20, 25, 26). The ABCF family members are unique in their structure and function as they lack transmembrane regions and therefore lack capacity for transport of substrates (24, 27). Of the ABCF family members, ABCF1 is most extensively characterized in eukaryotes, with functions ranging from initiation of mRNA translation, immune modulation, and nucleic acid sensing (27–32). The diverse functions attributed to ABCF1 are physiologically important, as demonstrated by the embryonic lethality of homozygous deletion of ABCF1 in mice(33). To date, nucleic acid sensing by ABCF1 has been defined using the dsDNA immunostimulatory DNA (ISD) sequence derived from *Listeria monocytogenes* (34) and a dsDNA HIV sequence, with both nucleic acid motifs inducing CXCL10, interferon-β expression, and downstream type I interferon responses in mouse embryonic fibroblasts(31). Complementary to dsDNA sensing, immune modulation mediated by ubiquitin-conjugating activities of ABCF1 have been defined in the context of macrophage polarization and immune responses linked to interferon-β production and tolerance important in mouse models of sepsis(32). In the context of studies using human lung samples, *ABCF1* gene expression has been identified in the human airway epithelium(35), although confirmation of protein and function remains to be determined. The clear *in vivo* demonstration of ABCF1 functions in immune responses in mouse models and the presence of detectable *ABCF1* gene expression the human airways warrants a deeper interrogation into the expression and function of this molecule in human health and disease.

Defining defence mechanisms in airway epithelial cells has important consequences in both lung health and disease, with the potential for interventions that could reduce viral-induced pathologies and exacerbations of chronic respiratory diseases(5–9). We therefore tested the hypothesis that ABCF1 functions as a dsDNA nucleic acid sensor in human airway epithelial cells important in regulating antiviral responses, using archived human lung samples and human airway epithelial cells. Expression and localization experiments were performed using *in situ* hybridization and immunohistochemistry in human lung tissue from healthy subjects, while confirmatory transcript and protein expression was performed in human airway epithelial cells. Functional experiments were performed with siRNA methods as no selective small molecule inhibitors to ABCF1 have been validated to date. Complementary transcriptomic analyses were performed to explore the potential contributions of ABCF1 beyond dsDNA virus sensing. Our results confirm expression of ABCF1 in human airway epithelial cells with a role for mediating CXCL10 production in response to dsDNA viral mimic challenge. Although, ABCF1 knockdown was associated with an attenuation of select genes involved in the antiviral responses, Gene Ontology analyses revealed a greater interaction of ABCF1 with TLR signaling suggesting a multifactorial role for ABCF1 in innate immunity in human airway epithelial cells.

## Methods

### Human Ethics

All studies using primary human lung material were approved by Hamilton integrated Research Ethics Board (HiREB – 5305-T and 5099-T).

### Reagents

*In situ* hybridization was performed using a custom RNAscope™ probe for ABCF1 (construct targeting 1713-2726 of NM_001025091.1) generated by Advanced Cell Diagnostics (ACD, Newark, California). Negative and positive control probes for quality control of RNA signal in analyzed human tissues were purchased from ACD (data not shown). Protein cell lysates were collected by lysing and scraping cells with RIPA Lysis buffer (VWR, Mississauga, Ontario) mixed with protease inhibitor cocktail (Sigma-Aldrich, Oakville, Ontario). Immunoblots were conducted using Mini-Protean TGX stain-free gels and Transfer-Blot Turbo RTA Transfer Kit reagents(Bio-Rad, Mississauga, Ontario). ABCF1 protein was probed with primary anti-ABCF1 antibody (HPA017578, Sigma-Aldrich, Oakville, Ontario) at 1:100 in 3% Casein in 1X Tris Buffered Saline with TWEEN ® 20 (Sigma-Aldrich, Oakville, Ontario, and Anti-rabbit HRP-linked Antibody (7074S - Cell Signaling Technology, Danvers, MA) at 1:2000. Immunohistochemistry was performed using the same anti-ABCF1 antibody as immunoblotting. ABCF1 and scramble siRNA SMARTpool siGENOME transfection reagents were purchased from Dharmacon (M-008263-01 and D-001206-14, Lafayette, Colorado). Cell viability was estimated with the Pierce LDH Cytotoxicity Assay kit (ThermoFisher Scientific, Mississauga, Ontario). RNA samples were lysed with Buffer RLT and purified with RNeasy Mini Kit columns (Qiagen, Toronto, Ontario). The ligands ISD, ISD control, VACV-70, VACV-70 control, and Poly:IC were all complexed with LyoVec transfection reagent and purchased from Invivogen (San Diego, California). Human CXCL10 was quantified using a commercial ELISA with ancillary reagent kit (R&D Systems, Oakville, Ontario). The protocol for quantifying CXCL10 was modified with the use of a loading plate for the samples.

### Cell culture

All experiments were performed in submerged monolayer cell culture. An immortalized human airway epithelial cell line (HBEC-6KT) over expressing human telomerase reverse transcriptase (hTERT) and cyclin-dependent kinase 4 (Cdk4) was used as previously described (36–40). HBEC-6KT were grown in keratinocyte serum free medium (ThermoFisher Scientific, Mississauga, Ontario) supplemented with 0.8 ng/mL epithelial growth factor, 50 μg/mL bovine pituitary extract and 1X penicillin/streptomycin. Calu-3 cells (ATCC HTB-55) were grown in Eagle’s Minimum Essential Media supplemented with 10% fetal bovine serum (Wisent, Saint-Jean-Baptiste, QC), 1mM HEPES, and 1X penicillin/streptomycin (Sigma-Aldrich, Oakville, Ontario). Primary human bronchial epithelial cells derived from healthy patient bronchial brushings were grown in PneumaCult ExPlus Medium supplemented with 96 μg/mL hydrocortisone (StemCell Technologies, Vancouver, BC) and 1X antimicrobial-antimycotics (ThermoFisher Scientific, Mississauga, Ontario). All cells were grown at 37*°C* at 5% CO_2._ Experiments with primary cells were performed between passages 1 and 4, and experiments with HBEC-6KT and Calu-3 cells were performed within 5 passages.

### *In vitro* experiments

All *in vitro* knockdown experiments in HBEC-6KT were done using siRNA transfected with DharmaFECT Transfection Reagent according to the manufacturer’s instructions. Cells were transfected with siABCF1 or siCTRL for 24 hours. After knockdown, cells were transfected with an immunostimulatory ligand for 24h followed by outcome measurements of cell viability (LDH and cell morphology), function (CXCL10 secretion), protein expression (immunoblot), or gene transcription (microarray). For TNF-α stimulation experiments, 10ng/ml was incubated for 24h followed by protein collection for immunoblots. For ISD and VACV-70 stimulation experiments, a concentration-response study was performed using 0.316-3.16μg/ml (ISD) or 0.1-3.16 μg/ml (VACV-70) followed by incubation for 24h. For Poly I:C stimulation experiments, 1μg/ml was incubated for 24h.

### Histology, digital slide scanning and microscopy

*In situ* hybridization and immunohistochemistry was performed using a Leica Bond Rx autostainer with instrument and application specific reagent kits (Richmond Hill, Ontario). The human lung tissues selected for analysis were formalin fixed, paraffin embedded, lung samples from archived hospital clinical samples, determined to be free of defined lung pathology. Following selection, four micron thick sections were stained with using RNAscope™ probes (*in situ* hybridization) or antibody (immunohistochemistry) following directions supplied with the Leica Bond reagent kits. For IHC, heat-induced antigen retrieval in citrate buffer was performed at pH 6 with primary antibody diluted at 1:50. Slides underwent digital slide scanning using an Olympus VS120-L100 Virtual Slide System at 40X magnification with VS-ASW-L100 V2.9 software and a VC50 colour camera (Richmond Hill, Ontario). Image acquisition and formatting was performed using Halo Software (Indica Labs, Albuquerque, NM).

### Gene Expression Omnibus (GEO) dataset mining

Gene expression patterns of *ABCF1* in human airway epithelial cells was determined relative to markers for immune cells (*CD34*), ABC transporters of known function in airway epithelial cells (*ABCC4*, *ABCC7*), and junctions (*CDH1*) in a dataset containing samples from trachea, large airways (generation 2^nd^-3^rd^), and small airways (generation 10^th^-12^th^) from healthy subjects (GSE11906, Affymetrix Human Genome U133 Plus 2 microarray platform)(41). The following probesets were used to extract gene expression data: *ABCF1* (200045_at), *ABCC4* (203196_at), ABCC7 (*CFTR*; 205043_at), *CDH1* (201131_s_at), and *CD34* (209543_s_at). In cases where more than one probe corresponded to a given gene, the following hierarchy was used to select an individual probe for further use: perfect, unique matches (probes ending in _at or _a_at) were preferred over mismatch or non-unique probes (ending in _s_at or _x_at). GSE11906 included 17 trachea (age – 42 +/− 7), 21 large airway (age – 42 +/− 9), and 35 small airway samples.

### Processing of raw microarray data

Raw intensity values from a microarray experiment using the Affymetrix Clariom S Human chip-type were imported into the R statistical language environment (version 3.6.1; R Core Team, 2019). Probe definition files were obtained from the Brainarray database (version 24(42)). The Single Channel Array Normalization (SCAN) method was used to obtain log_2_-transformed normalised expression values with the SCAN.UPC R package (version 2.26.0(43)), with annotation data from the Bioconductor project (version 3.9(44).

### Analysis of processed microarray data

From the processed expression values, principle component analyses were performed with the prcomp function (version 0.1.0) from the R statistical language (version 3.6.1; R Core Team, 2019) using default parameters. Determination of statistically significant differential gene expression was performed using the empirical Bayes method via the eBayes function from the limma R package (version 3.40.0(45). *P* values were adjusted using the Benjamini & Hochberg method, with a significance cutoff of 0.05. Significantly enriched Gene Ontology (GO) Biological Process Terms (ranked by *p* value) were determined using Enrichr ((46, 47)). Scatter plots, PCA plots, and GO term enrichment dot plots were generated using the ggplot2 R package (version 3.2.1). Heat maps were generated using the pheatmap R package (version 1.0.12), with log_2_ expression scaled by gene and complete hierarchical clustering using a Euclidean distance measure applied. A GO term enrichment clustergram was modified from Enrichr using Inkscape.

### Statistical analyses

All experiments were performed with an n≥3 unless otherwise noted. Experiments with HBEC-6KT and Calu-3 cells were considered independent when separated by a passage. Statistics were determined by permutation ANOVA with a Bonferonni-corrected post-hoc test comparing selected groups with *p*<0.05 determined to be statistically significant.

## Results

### ABCF1 gene and protein expression is localized to human airway epithelial cells *in situ* and *in vitro*

Expression and functional studies of ABCF1 have focused on human synoviocytes, mouse embryonic fibroblasts, human embryonic kidney cells, and peripheral blood mononuclear cells(27–30, 32). We have demonstrated gene expression of ABCF1 in human airway epithelial cells(35). To date, no *in situ* gene and protein expression data has confirmed ABCF1 expression in human lung tissues. To address this knowledge gap, we first mined publicly available gene expression data from primary human airway epithelial cells from healthy subjects. *ABCF1* gene expression was observed along the airway generations (trachea, large, and small) at levels relative to *ABCC7/CFTR* and *ABCC4*, two other ABC transporters with reported functions in airway epithelial cells (25, 40, 48, 49) (**Figure 1A**). CD34 and CDH1 (encoding E-Cadherin) were used as negative and positive control genes, respectively, for airway epithelial cells to provide contextual expression levels.

**Figure 1.**
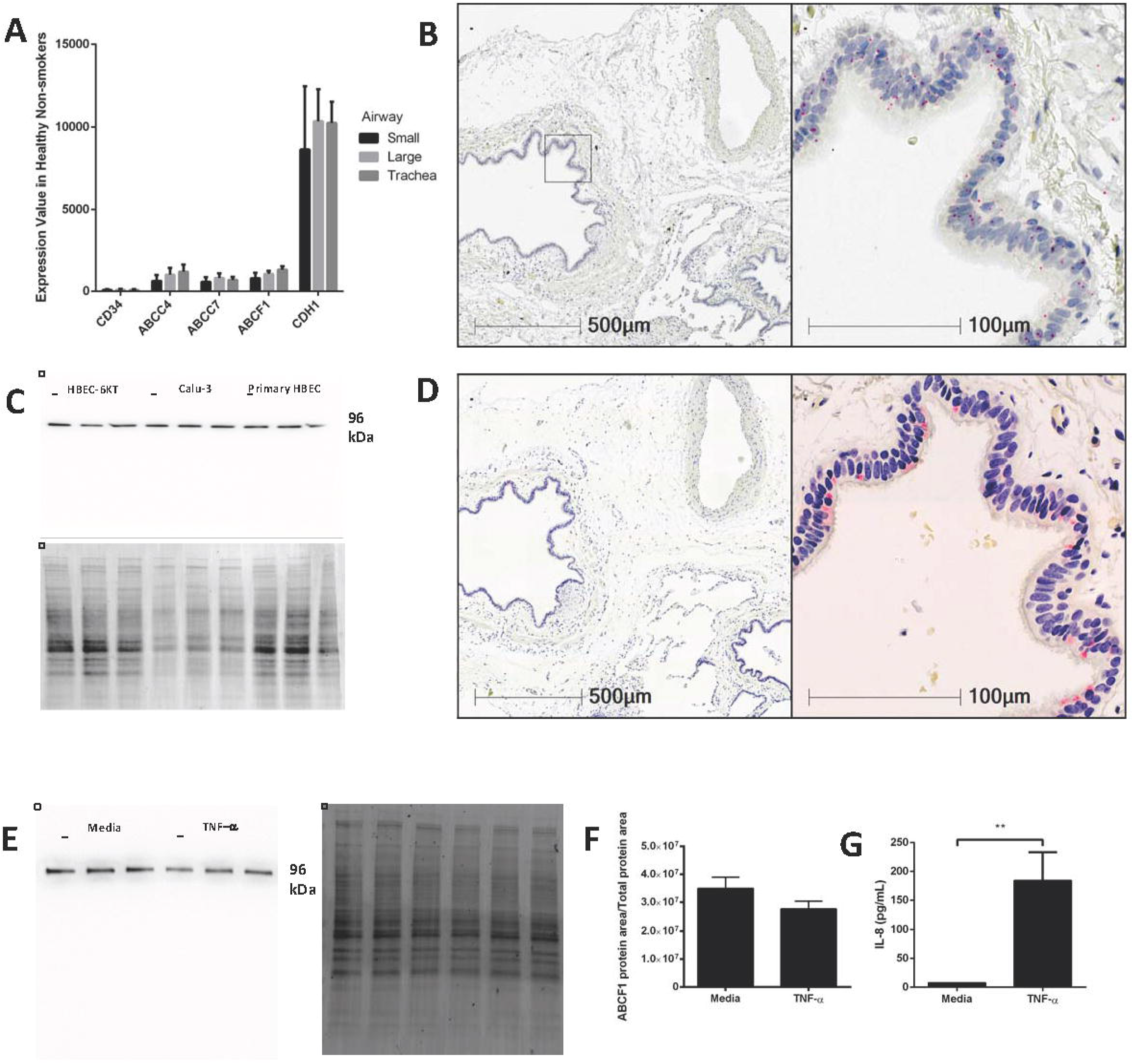
Validation of ABCF1 gene and protein expression in human airway epithelial cells i*n situ* and *in vitro*. **A**: Gene expression analysis of GEO deposited microarray dataset (GSE11906) generated from epithelial cells isolated from trachea, large (2^nd^-3^rd^ generation), and small airways (10^th^-12^th^ generation) from healthy subjects (see Methods for details). **B**: *In situ* hybridization of ABCF1 RNAscope™ probe in human lung under low (10X) and high (40X) magnification. Red puncta are representative of *ABCF1* gene transcripts with nuclei counterstained blue. Representative image of n=10. **C**: Immunoblot confirmation of ABCF1 protein expression in HBEC-6KT, Calu-3, and primary human airway epithelial cells (each cell type n=3 distinct cell line passages or donors) with a single band observed at predicted molecular weight (96kDa) with total protein loading blot demonstrating equal protein loading for each cell type. **D**: Immunohistochemistry of ABCF1 in human lung under low (10X) and high (40X) magnification. Representative image of n=10.Pink/red staining is representative of ABCF1 protein with nuclei counterstained blue. **E:** Immunoblot of ABCF1 following TNF-α stimulation of HBEC-6KT cells with corresponding total protein stain. **F:** Quantification of immunoblot of ABCF1 protein expression. **G:** IL-8 secretion from HBEC-6KT cells measured by ELISA as positive control for TNF-α stimulation. All studies n=3 unless otherwise notes. *=p<0.05

Next, *in situ* localization of ABCF1 gene transcript was performed using RNAscope™ probes on archived formalin fixed paraffin embedded human lung samples (**Figure 1B**). *ABCF1* gene transcript was observed in small puncta throughout the cytoplasm and nuclear areas of airway epithelial cells. ABCF1 staining was also observed in submucosal cells with morphology consistent with macrophages. Protein expression levels were next explored with validation of a commercially available antibody for ABCF1 using *in vitro* culture of primary human airway epithelial cells and two distinct airway epithelial cell lines (**Figure 1C**). For each airway epithelial cell type, a single band was observed at the predicted molecular weight of 96kDa for ABCF1, validating the use of the antibody for *in situ* immunohistochemistry localization. Consistent with *in vitro* ABCF1 protein expression data, positive staining was observed in human airway epithelial cells as shown in a serial section used for *in situ* hybridization (**Figure 1D**) with sparse staining in immune cells with macrophage morphology. Lastly, to explore proposed regulatory mechanisms for ABCF1(27), we performed a TNF-α exposure in human airway epithelial cells. Exposure to 10ng/ml TNF-α for 24h failed to induce a change in ABCF1 protein expression (**Figure 1E-F**), despite inducing an increase in IL-8 (**Figure 1G**). Collectively our *in vitro* and *in situ* data confirm gene and protein expression of ABCF1 in human airway epithelial cells, warranting downstream characterization and functional studies.

### *In vitro* attenuation of ABCF1 under basal conditions has limited impact on cell viability and transcriptional profiles

Functional studies have implicated ABCF1 in translation initiation and have demonstrated that homozygous loss of function results in embryonic lethality(28–30, 33). We therefore first interrogated the basal functions of ABCF1 in our human airway epithelial cells in the context of cell viability and transcriptional profiling.

We performed siRNA experiments to attenuate ABCF1 expression levels as no small molecule ABCF1 inhibitor has been described to date. Using siRNA approaches in human airway epithelial cells, we confirm that ABCF1 protein can be attenuated with qualitative (**Figure 2A**) and quantitative measures (**Figure 2B**). LDH levels were not elevated when ABCF1 was attenuated with siRNA **(Figure 2C**). Cell morphology was not different in ABCF1 attenuated human airway epithelial cells (**Figure 2D**). Collectively, the quantitative and qualitative data suggest moderate levels of siRNA knockdown are not associated with compromised human airway epithelial cell viability.

**Figure 2.**
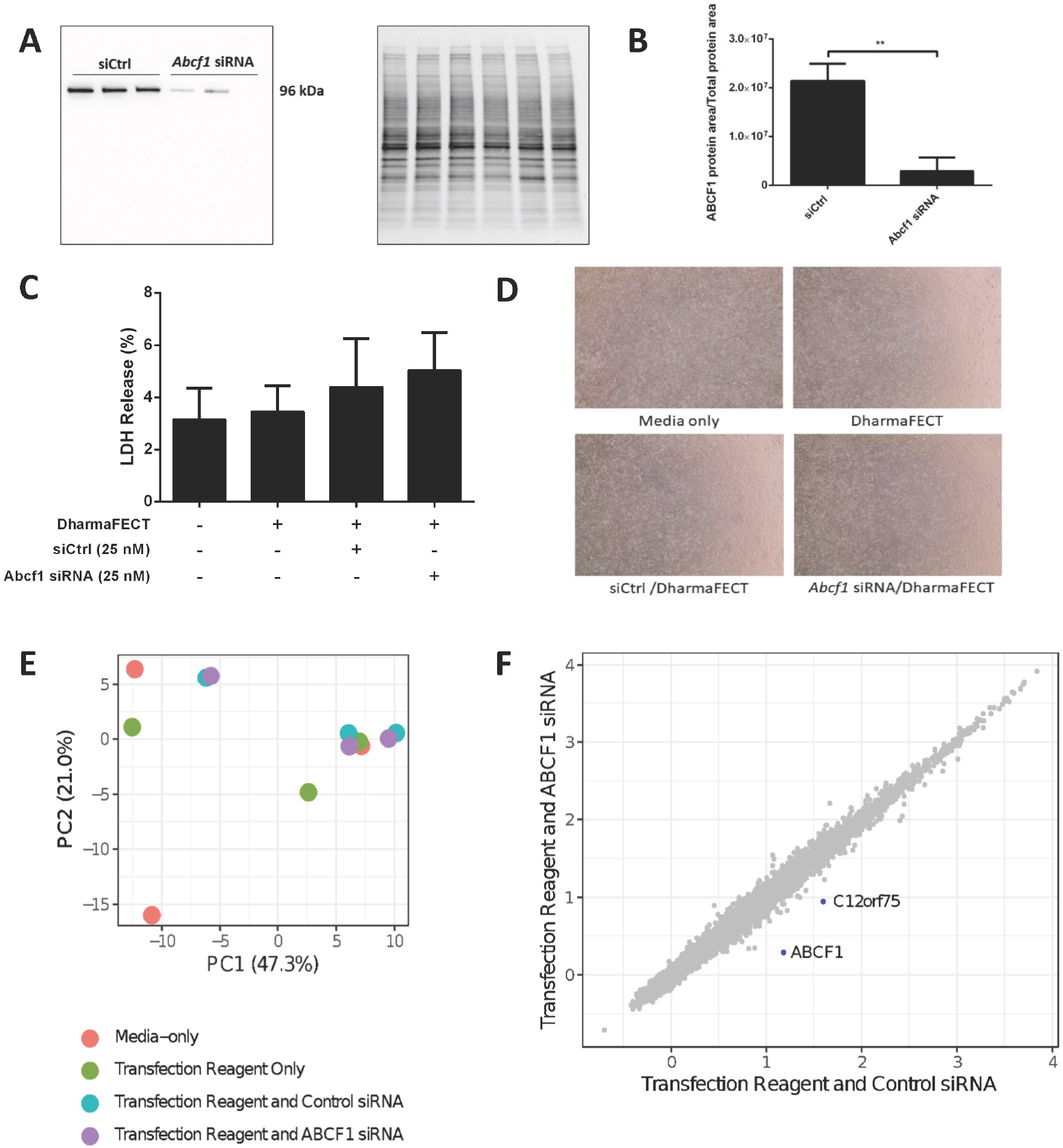
Interrogation of ABCF1 under basal conditions in human airway epithelial cells *in vitro*. **A**: Immunoblot confirming siRNA-mediated knockdown of ABCF1 protein expression in HBEC-6KT cells. **B**: Quantification of ABCF1 protein expression following siRNA treatment. **C**: LDH quantification as a measure of cell viability following siRNA treatment. **D**: Phase-contrast microscopy (4X magnification) of HBEC-6KT following siRNA treatment. **E**: PCA plot of microarray gene expression profiles of HBEC-6KT cells following siRNA treatment. Red circles (media alone), green circles (transfection reagent only), blue circles (transfection reagent and control siRNA), purple circles (transfection reagent and ABCF1 siRNA). **F**: Log_2_ expression data for transfection reagent with control siRNA compared to transfection reagent with ABCF1 siRNA. Significantly differently expressed genes are in blue and are down-regulated (*ABCF1* and *C12orf75*). All studies n=3. * = *p*<0.05.

To interrogate the impact of ABCF1 attenuation under basal conditions, a human gene expression microarray analysis was performed. A principal component analysis of ABCF1 attenuation and corresponding experimental controls revealed no clustering between experimental replicates for any condition (**Figure 2E**), suggesting that the overall impact of ABCF1 attenuation under basal conditions minimally impacted global gene expression patterns. Statistical analysis comparing ABCF1 attenuation and siRNA control treated human airway epithelial cells confirmed *ABCF1* gene was down-regulated (**Figure 2F**) which was associated with only one other significantly differentially expressed (up or down) gene, *C12orf75*, which encodes overexpressed in colon carcinoma-1 (OCC-1) protein.

Collectively our *in vitro* studies under basal conditions demonstrate that ABCF1 attenuation is not associated with changes in viability or significant genome wide changes in transcriptional profiles in human airway epithelial cells.

### The dsDNA viral mimic VACV-70 induces CXCL10 and an antiviral response in human airway epithelial cells *in vitro*

As attenuation of ABCF1 under basal conditions resulted in limited impacts on cell viability and gene transcription, we next explored conditions of *challenge* in human airway epithelial cells. ABCF1 was described as a dsDNA sensor in mouse embryonic fibroblasts that mediated CXCL10 secretion under challenge conditions with the viral mimic interferon stimulatory DNA (ISD) sequence(31), a 45bp oligomer shown to activate the STING-TBK1-IRF3 antiviral sensing axis (34, 50).

To determine the response of human airway epithelial cells to ISD, we performed a concentration-response study followed by quantification of extracellular CXCL10 secretion (**Figure 3A**). ISD induced an increase in CXCL10 at 1ug/ml while trends were observed at lower (0.316ug/ml) and higher (3.16ug/ml) concentrations. Importantly, as concentration of ISD increased, the cellular response to the control (ssDNA of the ISD sequence) also increased. These results limited the use of ISD as dsDNA challenge stimulus in human airway epithelial cells for studying ABCF1 function.

**Figure 3.**
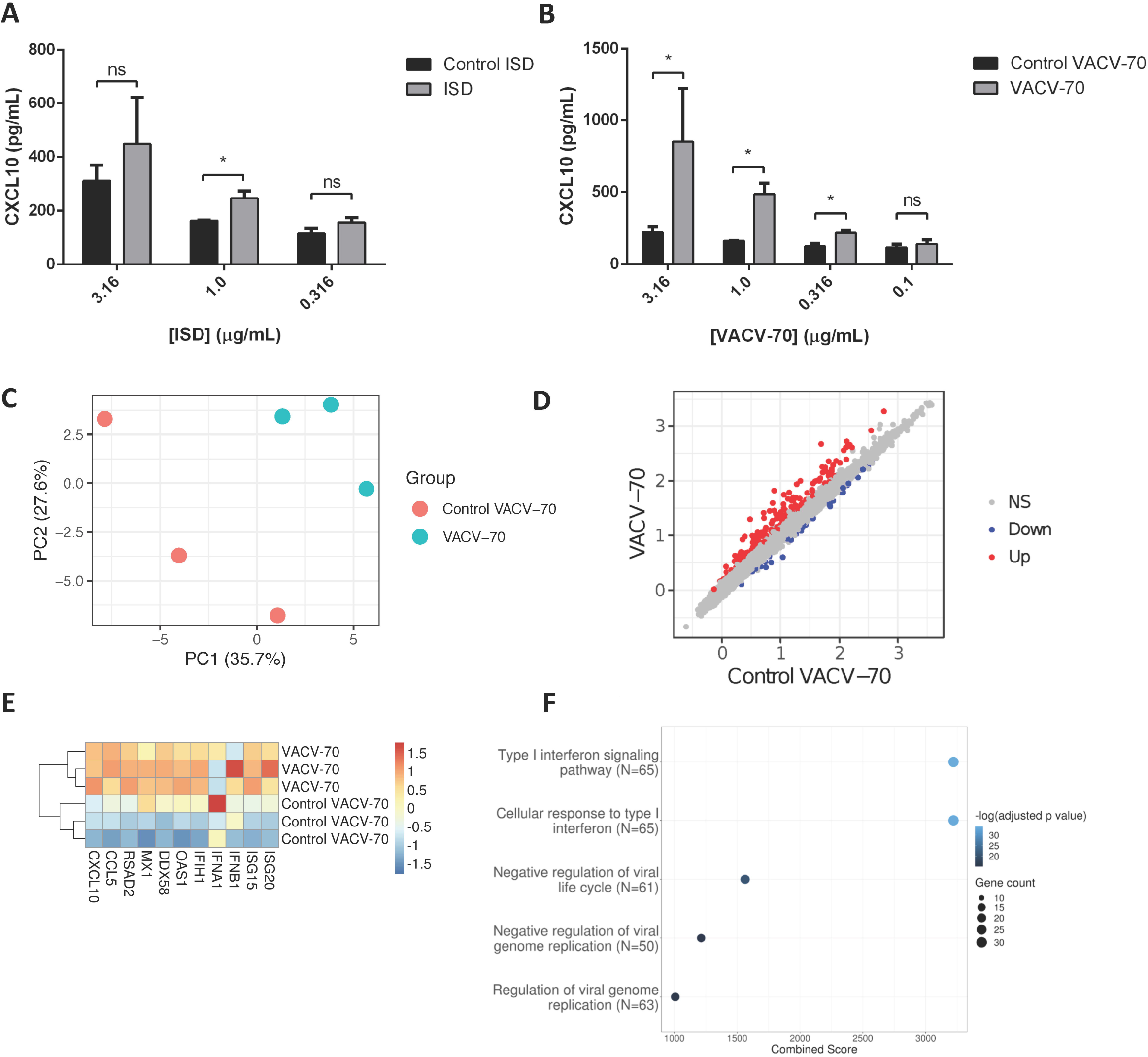
dsDNA induced antiviral responses in human airway epithelial cells *in vitro.* **A**: Concentration-response analysis of ISD-induced CXCL10 protein production for HBEC-6KT cells. Grey bars: ISD, Black bars: control ssDNA generated from ISD. **B**: Concentration-response analysis of VACV-70-induced CXCL10 protein production for HBEC-6KT cells. Grey bars: VACV-70, Black bars: control ssDNA generated from VACV-70. **C**: PCA plot of microarray gene expression profiles of HBEC-6KT cells following VACV-70 or control. Red circles (control VACV-70), blue circles (VACV-70). **D**: Log_2_ expression data for transfection treatment with control VACV-70 compared to VACV-70. Significantly differently expressed genes are identified in blue (down – 42 genes) and red (up – 170 genes). **E**: Heat map of log_2_ expression data of select known antiviral genes scaled by gene were extracted for VACV-70 and Control VACV-70 samples. **F**: Top 5 GO Biological Processes are ranked by increasing −log_10_ adjusted *p* value, with number (Count) of significantly differentially expressed genes between VACV-70 and control VACV-70 contributing to the total number of genes associated with the given pathway (N) denoted by the size of circle. All studies n=3. * = *p*<0.05.

Vaccinia virus is a dsDNA virus that is able to infect airway epithelial cells(19–23). We therefore determined the response of human airway epithelial cells to VACV-70, a 70bp dsDNA oligonucleotide containing Vaccinia virus motifs(51). VACV-70 induced a concentration dependent increase in CXCL10 from 0.316ug/ml to 3.16ug/ml. In contrast to ISD, no cellular response to the control (ssDNA of the VACV-70 sequence) was observed at any concentration.

To characterize the molecular pathways activated by VACV-70, we performed a transcriptional and pathway analysis of human airway epithelial cells following challenge. To interrogate the VACV-70 transcriptional responses a principal component analysis was performed for microarray gene expression data, revealing distinct clustering between stimulation (VACV-70) and control (**Figure 3C**). Statistical analysis revealed 170 up-regulated genes and 42 down-regulated genes with VACV-70 stimulus (**Figure 3D**). VACV-70 up-regulated *CXCL10* gene expression and a curated list of antiviral related interferon stimulated genes (**Figure 3E)**. GO term analysis revealed that the top pathways activated by VACV-70 were associated with type I interferon signaling, viral responses, and cellular responses to viruses (**Figure 3F**).

Collectively our *in vitro* challenge studies confirm that VACV-70, a dsDNA viral mimic, can induce CXCL10 and antiviral transcriptional responses in human airway epithelial cells.

### *In vitro* attenuation of ABCF1 under VACV-70 stimulated conditions attenuates CXCL10 secretion with limited impact on cell viability

We have confirmed VACV-70 induction of CXCL10 in human airway epithelial cells at the gene **(Figure 3E**) and protein (**Figure 3B**) levels. Furthermore, we have demonstrated siRNA-mediated attenuation of ABCF1 with no impact on cell viability(**Figure 2A-D**). We therefore performed a VACV-70 challenge with ABCF1 attenuation by siRNA knockdown with a readout of CXCL10.

ABCF1 knockdown was associated with a decrease in CXCL10 protein secretion from human airway epithelial cells, with confirmation and quantification of ABCF1 knockdown performed by immunoblot (**Figure 4A-C**). Cell viability following VACV-70 challenge and ABCF1 attenuation was not impacted as assessed by LDH quantification (**Figure 4D**). Qualitative analysis following VACV-70 challenge and ABCF1 revealed no impact on human airway epithelial cell of cell morphology (**Figure 4E**).

**Figure 4.**
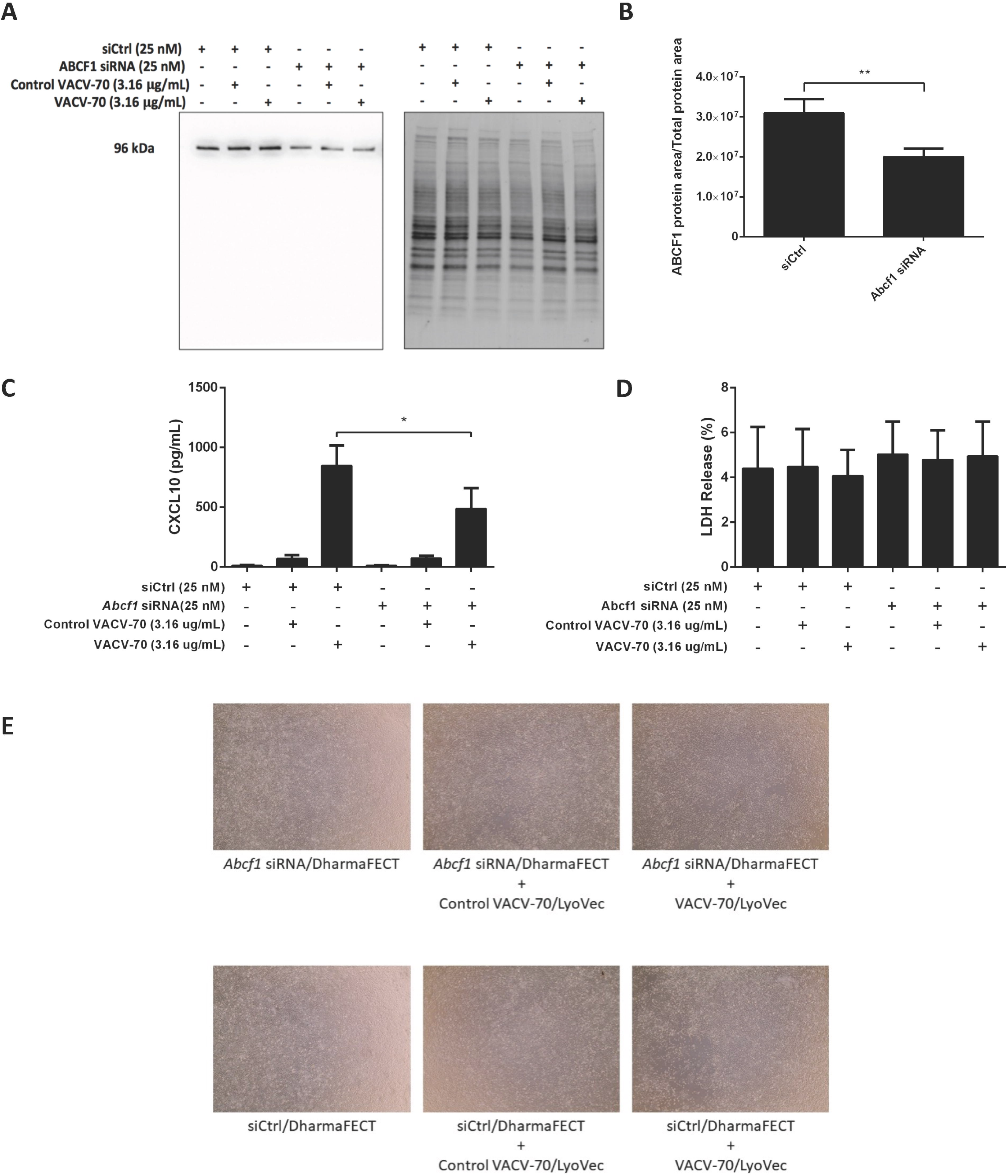
ABCF1 knockdown attenuates VACV-70 induced CXCL10 production from in human airway epithelial cells *in vitro*. **A**: Immunoblot confirming siRNA-mediated knockdown of ABCF1 protein expression in HBEC-6KT cells under experimental conditions of VACV-70 challenge **B**: Quantification of ABCF1 protein expression following siRNA treatment. **C**: VACV-70 (3.16ug/ml)-induced CXCL10 protein production for HBEC-6KT cells with ABCF1 siRNA treatment. Black bars: siCtrl treated. Grey bars: ABCF1 siRNA treated. **D:** LDH quantification as a measure of cell viability following VACV-70 and siRNA treatment. **E**: Phase-contrast microscopy (4X magnification) of HBEC-6KT following VACV-70 and siRNA treatment. All studies n=3. * = p<0.05.

Collectively our *in vitro* challenge and functional studies demonstrate that ABCF1 siRNA treatment attenuated VACV-70-induced CXCL10 protein secretion in human airway epithelial cells.

### ABCF1 attenuation does not impact VACV-70-induced antiviral transcriptional responses in human airway epithelial cells

In parallel to induction of *CXCL10* gene, we have confirmed with GO pathway analysis that VACV-70 induces a dominant antiviral transcriptional signature (**Figure 3E-F**). We therefore next explored how ABCF1 attenuation impacts transcriptional responses induced by VACV-70, beyond induction of CXCL10.

A principal component analysis was performed for microarray gene expression data, revealing distinct clustering in samples where ABCF1 was attenuated relative to control conditions under conditions of VACV-70 challenge (**Figure 5A – green vs purple)**. Statistical analysis revealed 63 up-regulated genes and 65 down-regulated genes when comparing ABCF1 attenuation relative to control under conditions of VACV-70 challenge (**Figure 5B**). siRNA knockdown of ABCF1 was confirmed and associated with attenuation of *CXCL10* gene expression(**Figure 5C**, p=0.06).

**Figure 5.**
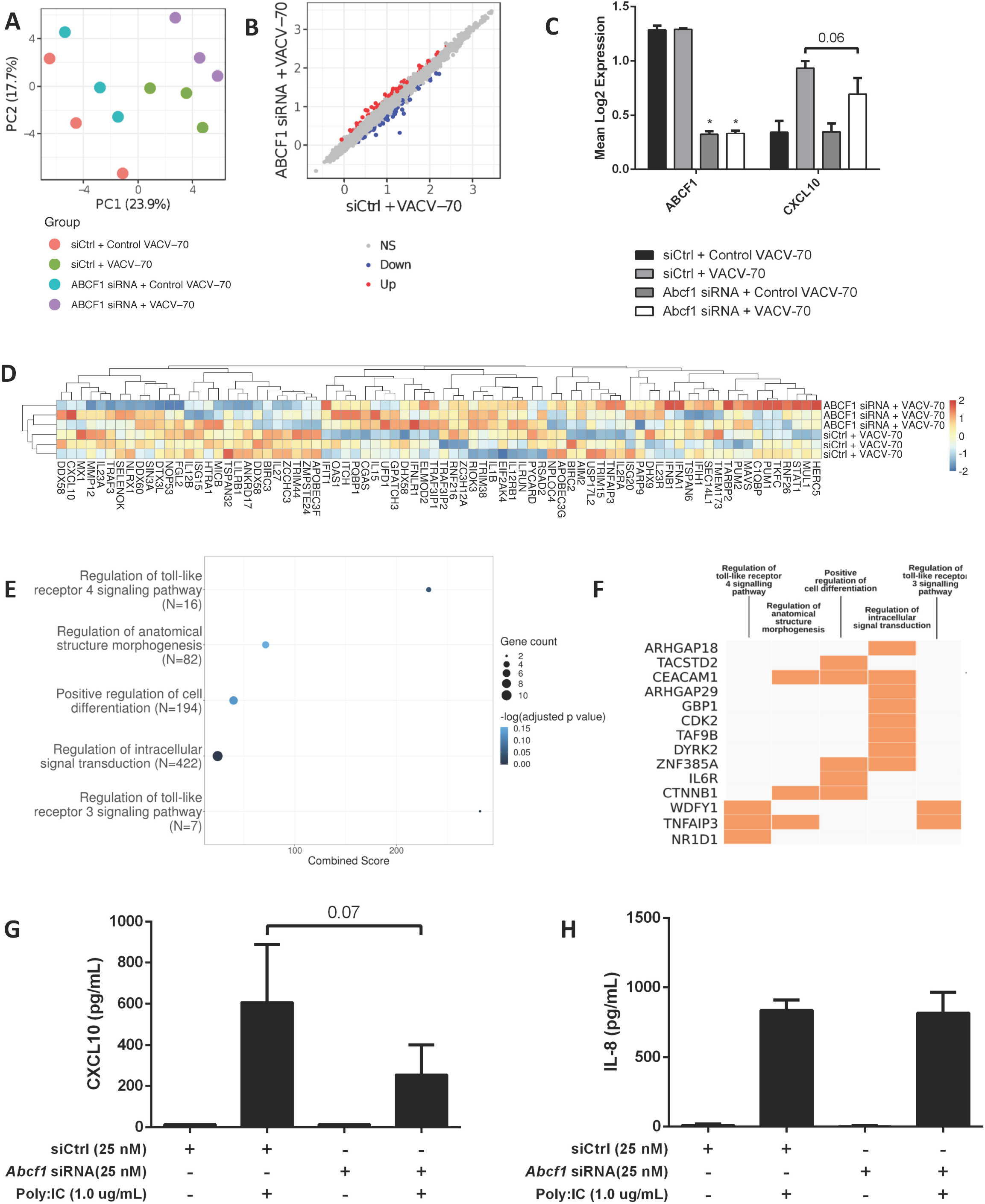
Transcriptional interrogation of ABCF1 siRNA knockdown and VACV-70 challenge in human airway epithelial cells *in vitro*. **A**: PCA plot of microarray gene expression profiles of HBEC-6KT cells following ABCF1 knockdown and VACV-70 treatment. Red circles (control siRNA and control VACV-70), green circles (control siRNA and VACV-70), blue circles (ABCF1 siRNA and control VACV-70), purple circles (ABCF1 siRNA and VACV-70). **B**: Log_2_ expression data for transfection treatment with control siRNA and VACV-70 compared to ABCF1 siRNA and VACV-70. Significantly differently expressed genes are identified in blue (down – 65 genes) and red (up – 63 genes). **C**: Confirmation of ABCF1 and CXCL10 attenuation with ABCF1 siRNA treatment presented as log_2_ expression data. Black bars: control siRNA and control VACV-70, light grey bars: control siRNA and VACV-70, dark grey bars: ABCF1 siRNA and control VACV-70, white bars: ABCF1 siRNA and VACV-70. **D:** Heat map of log_2_ expression data of genes associated with the regulation of defense response to virus GO term (n=68) plus the selected known antiviral genes from Figure 3E (n=11), scaled by gene were extracted for VACV-70 samples with and without ABCF1 siRNA. **E**: Top 5 GO Biological Processes are ranked by increasing −log_10_ adjusted *p* value, with number (Count) of significantly differentially expressed genes between VACV-70 samples with and without ABCF1 siRNA, contributing to the total number of genes associated with the given pathway (N) denoted by the size of circle. **F**: Significantly differentially expressed genes between VACV-70 samples with and without ABCF1 were submitted to Enrichr for generation of a clustergram defining the gene contribution (Y-axis) to the functional enrichment of the top 5 GO Biological Processes (X axis), with orange squares denoting the association of a differentially expressed gene with a particular GO term. **G:** Poly I:C (1.0 ug/ml)-induced CXCL10 and **H:** IL-8 secretion for HBEC-6KT cells with ABCF1 siRNA treatment. All studies n=3. * = p<0.05.

To explore a focused transcriptional response of ABCF1 attenuation in the context of VACV-70 challenge, a hypothesis-directed approach curated 79 genes from the GO term “Regulation of defense response to virus” and key components of viral sensing for heat map visualization (52)(**Figure 5D**). Statistical analysis revealed no global significant difference between ABCF1 attenuated and control groups for the expression pattern of this curated list of genes.

To explore the broader transcriptional responses of ABCF1 attenuation in the context of VACV-70 challenge, a hypothesis-free directed approach with GO term analysis was performed. Top-ranking GO pathway terms included *Regulation of toll-like receptor 3-4 signaling pathways*, which were driven by the genes *WDFY1*, *TNFAIP3*, and *NR1D1* (**Figure 5E-F**).

As our data suggested that ABCF1 functions in human airway epithelial cells may extend beyond sensing of VACV-70 dsDNA viral mimic through regulation of TLR signaling, we explored Poly I:C, a dsRNA analog and TLR3 ligand that induces interferon responses including CXCL10 production. ABCF1 attenuation was associated with a trend (p=0.07) for reduced Poly I:C-induced CXCL10 protein secretion but did not impact Poly I:C-induced IL-8 protein secretion (**Figure 5G-H**). Collectively our *in vitro* challenge and functional studies with transcriptional analyses demonstrate a role for ABCF1 in mediating VACV-70 and Poly I:C induced CXCL10 secretion and TLR4 related signaling in human airway epithelial cells.

## Discussion

ABCF1, a member of the ATP Binding Cassette family expressed in diverse mammals and different tissue types, has been reported to have diverse functions including initiation of mRNA translation, dsDNA viral sensing, and polarization of immune cell phenotype(27–32). We have recently reported ABCF1 gene expression levels in human airway epithelium(35), but the function of this molecule remained unexplored. Herein we confirm ABCF1 gene and protein expression *in situ* and *in vitro* in primary human lung tissue and cell lines and explore its function in airway epithelial cells. Under basal conditions, ABCF1 attenuation did not lead to quantitative changes in cell viability or qualitative changes in cell morphology associated with cell death. Furthermore, ABCF1 attenuation failed to significantly alter basal transcriptional activity in human airway epithelial cells. Under VACV-70 challenge, a model of dsDNA viral exposure, ABCF1 was linked to CXCL10 secretion. Interestingly, despite the demonstrated activation of a viral gene signature by VACV-70, no global change in antiviral gene expression patterns were observed with ABCF1 attenuation. In contrast, the gene pathways regulated by ABCF1 under VACV-70 challenge were associated with TLR signaling and intracellular signal transduction. Furthermore, Poly I:C, a dsRNA analog and TLR3 ligand induced CXCL10 in an ABCF1 sensitive mechanism. Collectively, our data suggests that ABCF1 may regulate CXCL10 production downstream of dsDNA sensing mechanisms and TLR3 in human airway epithelial cells.

ABCF1 (originally called ABC50) was first identified in human synoviocytes at the mRNA level as a transcript regulated by TNF-α exposure(27). ABCF1 is unique in the mammalian ABC transporter family in that it contains the signature ATP binding LSGGQ amino acid motif and associated Walker A and B motifs for phosphate binding, but lacks a predicted transmembrane region(27, 53, 54), which is supportive of a cytosolic localization and function. *ABCF1* transcript expression profiling has revealed near ubiquitous expression in human organs including lung, heart, brain, placenta, liver, skeletal muscle, kidney, pancreas, spleen, thymus, prostate, testis, ovary, small intestine, colon, peripheral blood leukocytes(27). The expression of ABCF1 has subsequently been identified in the human HeLa cells and embryonic kidney cells and other mammalian cells from rats, rabbits, hamsters, and mice (28–30, 32, 33). Highlighting the importance of ABCF1 in normal physiology and development, homozygous deletion of ABCF1 is embryonic lethal in either C57Bl/6 mice or BALB/c mice(33). As our group recently identified gene expression of *ABCF1* in human airway epithelial cells (35), we set out to first confirm this at the protein level and then determine the function(s) of ABCF1 in human airway epithelial cells. We confirm that *ABCF1* gene expression is present in airway epithelial cells and expressed at levels relative to other known ABC transporters with function in this cell type, *ABCC4* and *ABCC7*/*CFTR*(25, 40, 48, 49). *In situ* hybridization using RNAscope™ technology demonstrated *ABCF1* transcripts present in the airway epithelial cells in healthy human lung samples, which was consistent with positive immunohistochemical staining of protein in a serial section of the same samples using an antibody validated for specificity. Since the original discovery of ABCF1 was the result of an upregulated transcript resulting from TNF-α stimulation of synoviocytes, we examined if this mechanism was functional in human airway epithelial cells. In contrast to the reported data on synoviocytes, TNF-α stimulation failed to induce ABCF1 protein expression in human airway epithelial cells, despite IL-8 induction as a positive control. The contrasting observations may be due to our measurement of ABCF1 protein whereas the original discovery focused on gene expression as a readout. Interestingly, in a recent report profiling the role of ABCF1 in murine bone-marrow derived macrophages, TNF-α stimulation *suppressed* ABCF1 protein expression(32). Collectively our results and those in the literature support gene and protein expression of ABCF1 in human airway epithelial cells, and that regulation of this protein is likely to be cell specific.

The first description of a potential function for ABCF1 in mammalian cells was derived from the experiments on human synoviocytes, suggesting a role in translation due to homology of molecular sequence with yeast proteins that performed this function(27, 28). The embryonic lethality observed in mice for homozygous ABCF1 deletion and ubiquitous expression across multiple cell and tissue types(33), is consistent with ABCF1 playing a role in a fundamental biological process like protein translation. The observation that proliferating cells including synoviocytes stimulated with TNF-α and T cells stimulated with phorbol myristate acetate and ionomycin, elevate ABCF1 levels is further consistent with a role in translation(27, 28). Subsequent to the discovery of ABCF1 gene expression and homology modeling, biochemical studies implicated the protein in interaction with eukaryotic initiation factor-2 (eIF2), a heterotrimeric protein consisting of α, β, and γ subunits, that is important for translation initiation(28). A distinguishing feature of ABCF1 relative to other ABC transporters is a N-terminal domain that is able to interact with eIF2α in a process that potentiates binding of methionyl-tRNA and initiation of translation(29). In addition to eIF2α interactions, ABCF1 associates with ribosomes in a process potentiated by ATP binding to the nucleotide binding domains and inhibited by ADP(28), although the hydrolysis of ATP seems dispensable for ribosome interaction(29). To explore the potential function of ABCF1 as an initiator of translation in human airway epithelial cells, we undertook a siRNA approach to attenuate gene and protein expression followed by a global assessment of cell viability and transcriptomics. Surprisingly, under basal conditions, ABCF1 attenuation at the gene and protein level did not impact human airway epithelial cell viability, morphology or transcriptional profile. Importantly, our outcome measurements were performed on human airway epithelial cells that were sub-confluent and undergoing proliferation in serum-free media, an experimental condition where ABCF1 function in translation initiation would be relevant. A limitation of our design is that we measured global gene expression under the assumption that this would reflect any global changes in gene translation, an indirect approach which does not allow us to directly implicate ABCF1 expression levels to protein synthesis. Interestingly, our observations of minimal changes in human airway epithelial cells may be consistent with cells of epithelial lineage, as near complete ABCF1 knockdown in HeLa cells was also only associated with a modest attenuation of total protein synthesis(30). Collectively, our results suggest that ABCF1 may function independent of protein translation functions in human airway epithelial cells, as gene and protein attenuation results in no changes in cell viability or global transcriptional profile.

The original discovery that ABCF1 expression was regulated by TNF-α stimulation suggested a link to immune responses, although no differential expression patterns were observed for synoviocytes from healthy individuals or those with rheumatoid arthritis(27). Subsequently, ABCF1 has been implicated in immune responses via a cytosolic dsDNA viral sensing function using mouse embryonic fibroblasts(31). Using an integrative bioinformatic and molecular biology approach, a biotinylated ISD sequence was used as a bait and transfected into cells, followed by proteomic interrogation of identified candidates. The ISD bait method was validated by identifying known dsDNA sensors including HMGB1, HMGB2, and HMGB3, components of the AIM2 inflammasome, and the SET complex that plays a role in HIV-1 retroviral detection and infection(55). Within the pool of unknown dsDNA interacting candidates, ABCF1 was mechanistically linked to ISD induced-CXCL10 production using siRNA methods. The observed ISD induced-CXCL10 converged on IRF3 signaling, confirmed by showing reduced IRF3 phosphorylation following ISD treatment under conditions of ABCF1 silencing. In a separate study, ABCF1 has been implicated as a molecular switch downstream of TLR4 signaling in mouse bone-marrow derived macrophages that regulates MyD88 dependent pro-inflammatory and TRIF/TRAM dependent anti-inflammatory processes(32). Using *in vitro* an *in vivo* model systems, ABCF1 was implicated in polarizing pro-inflammatory macrophages to an anti-inflammatory/tolerant macrophage phenotype with direct involvement in shifting the systemic inflammatory response syndrome to a endotoxin tolerance phase in sepsis(32). The mechanism responsible for the ABCF1-mediated polarization of macrophages was identified to be a E2-ubiqutin-conjugating enzyme function. In wild-type macrophages the TRIF-IFN-β pathway is intact with attenuation of the MyD88 pathway, allowing IRF-3 phosphorylation, dimerization, and IFN-β expression. In contrast, heterozygosity for ABCF1 results in attenuation of the TRIF-IFN-β pathway, with reduced IRF-3 activation and IFN-β production. Importantly, these two immunological studies converge on a relationship between ABCF1 and IRF3, which could involve direct or indirect interactions to facilitate downstream immune responses. Consistent with the potential role for ABCF1 as a dsDNA sensor, we explored immune and transcriptional responses downstream of VACV-70, a dsDNA viral mimic capable of activating STING, TBK1, and IRF3 independent of TLRs(51). VACV-70 induced a dominant antiviral signature and pathway activation in human airway epithelial cells, consistent with successful transfection and cytosolic sensing. ABCF1 attenuation was associated with a reduction in CXCL10, an antiviral cytokine regulated by IRF3 activation, independent of any changes in cell viability or morphology. Transcriptomics revealed that although attenuation of CXCL10 was observed with ABCF1 siRNA, a global attenuation of an antiviral signature was not observed. Hypothesis-free GO analysis identified that the key pathways that were significantly impacted by ABCF1 siRNA treatment during VACV-70 challenge were related to TLR signaling. Interestingly, a key gene identified in our VACV-70 challenge and ABCF1 attenuation studies is *WDFY1*, which links TLR3/4, TRIF, and IRF3 signaling(56). This finding suggested that ABCF1 could potentially be regulating both TLR4 and TLR3/TRIF/IRF3 signaling (32). We tested this hypothesis by using Poly I:C, a dsRNA viral mimic that activates TLR3 and IRF3 (52). ABCF1 siRNA treatment attenuated Poly I:C-induced CXCL10 production, further demonstrating a link between ABCF1 and TRIF/IRF3, perhaps through regulation of *WDYF1*. Our exploratory results suggest that ABCF1 is likely to play a complex role in innate immunity in response to cytosolic nucleic acids, with a potential interaction with TRIF/IRF3 for regulation of CXCL10.

Throughout our study we encountered several technical issues. The absence of pharmacological interventions that could antagonize ABCF1 function required us to pursue molecular approaches with siRNA. siRNA approaches were unable to completely attenuate ABCF1 at concentrations of 25nM for up to 48h. Longer durations of silencing were not possible as the human airway epithelial cell line used showed changes in morphology with vehicle control transfection reagent beyond 48h of incubation. Our inability to completely attenuate ABCF1 levels was consistent with human embryonic kidney cells(30). Secondary to addressing ABCF1 expression levels, we sought to explore the functional consequences with the reported dsDNA viral mimic ISD as reported in the literature with mouse embryonic fibroblasts(32). During our concentration-response studies with ISD, the vehicle control condition resulted in elevations in our primary readout of CXCL10, which suggested an unexplained confounding factor. We therefore opted to use VACV-70 in place of ISD, which limits our ability to directly compare our results to those that have established ABCF1 as a dsDNA sensor with ISD(32).

In conclusion, we confirm that ABCF1 is expressed at the gene and protein level *in situ* and *in vitro* in human airway epithelial cells. In human airway epithelial cells, ABCF1 has minimal functions for cell viability and transcriptional regulation under basal conditions, but is important for mediating immune responses to cytosolic nucleic acids in pathways that involve TLR signaling and CXCL10 production. Our data form the foundation to pursue precisely how ABCF1 is regulated and where it functions in the network of cytosolic nucleic acid sensors and immune responses in human airway epithelial cells.

## Author contributions

QC: Designed, performed, and analyzed *in vitro* experiments, *in vitro* figure generation, drafting and editing of the manuscript.

JA: Performed bioinformatics analysis, figure generation, and drafted the manuscript.

BT: Performed bioinformatics analysis, figure generation, and drafted the manuscript.

NA: Performed *in vitro* experiments, & *in vitro* figure generation.

NT: Processed human tissue, performed *in vitro* experiments, & *in vitro* figure generation

NM: Responsible for patient consenting and human tissue acquisition.

SR: Responsible for histology and imaging.

AA: Responsible for human ethics protocols, human tissue acquisition, histology, and imaging.

GC: Responsible for supervision of the clinical research coordinator and human tissue acquisition.

KA: Responsible for supervision of the trainees, human tissue acquisition, study design, histology, and imaging.

AD: Responsible for oversight of the entire study & supervision of the trainees and funding.

JH: Responsible for oversight of the entire study & supervision of the trainees and funding, drafting and editing of the manuscript.

